# The circadian clock regulates PIF3 protein stability in parallel to red light

**DOI:** 10.1101/2023.09.18.558326

**Authors:** Wei Liu, Harper Lowrey, Chun Chung Leung, Christopher Adamchek, Juan Du, Jiangman He, Meng Chen, Joshua M. Gendron

## Abstract

The circadian clock is an endogenous oscillator, but its importance lies in its ability to impart rhythmicity on downstream biological processes or outputs. Focus has been placed on understanding the core transcription factors of the circadian clock and how they connect to outputs through regulated gene transcription. However, far less is known about posttranslational mechanisms that tether clocks to output processes through protein regulation. Here, we identify a protein degradation mechanism that tethers the clock to photomorphogenic growth. By performing a reverse genetic screen, we identify a clock-regulated F-box type E3 ubiquitin ligase, *CLOCK-REGULATED F-BOX WITH A LONG HYPOCOTYL 1* (*CFH1*), that controls hypocotyl length. We then show that CFH1 functions in parallel to red light signaling to target the transcription factor PIF3 for degradation. This work demonstrates that the circadian clock is tethered to photomorphogenesis through the ubiquitin proteasome system and that PIF3 protein stability acts as a hub to integrate information from multiple environmental signals.

## Main text

The circadian clock coordinates biological processes to specific times of day. In plants, the core circadian clock is composed of transcriptional feedback loops, but 24-hour rhythmicity requires post-translational modification of transcription factors and eventually regulated degradation by the ubiquitin proteasome (*1–3*). The transcription factors that make up the core of the plant circadian clock also control the timing of output processes through direct regulation of gene transcription, but evidence suggests that there are rich hierarchical post-transcriptional and post-translational networks that tether the clock to outputs to ensure their 24-hour rhythmicity (*4, 5*). Efforts have led to increased understanding of the transcriptional connections between the circadian clock and its outputs, but we know far less about the post-translational connections, including degradation mechanisms that impart 24-hour rhythms on proteins that participate in these output processes.

To discover protein degradation mechanisms that tether the circadian clock to downstream biological processes, we identified F-box type E3 ubiquitin ligase genes whose expressions are controlled by the circadian clock. We searched published Arabidopsis microarray datasets for F-box genes with rhythmic expression in constant light (Fig. 1A) (*6*) and chose genes that were rhythmic in at least 2 of the 3 available experiments (correlation cutoff at 0.8) for further study. This included 31 F-box genes with 19 rhythmic in two of three experiments and 12 rhythmic in all three (Table S1, Fig. S1). Supporting our approach, two of the identified genes are known to tether the circadian clock to important biological processes. For example, *FLAVIN-BINDING KELCH REPEAT F-BOX 1* (*FKF1*) functions as a key clock-controlled flowering time regulator, and *PHLOEM PROTEIN 2-A13* (*PP2-A13*) is required for growth in winter photoperiods in Arabidopsis (*7–10*).

**Fig. 1.**
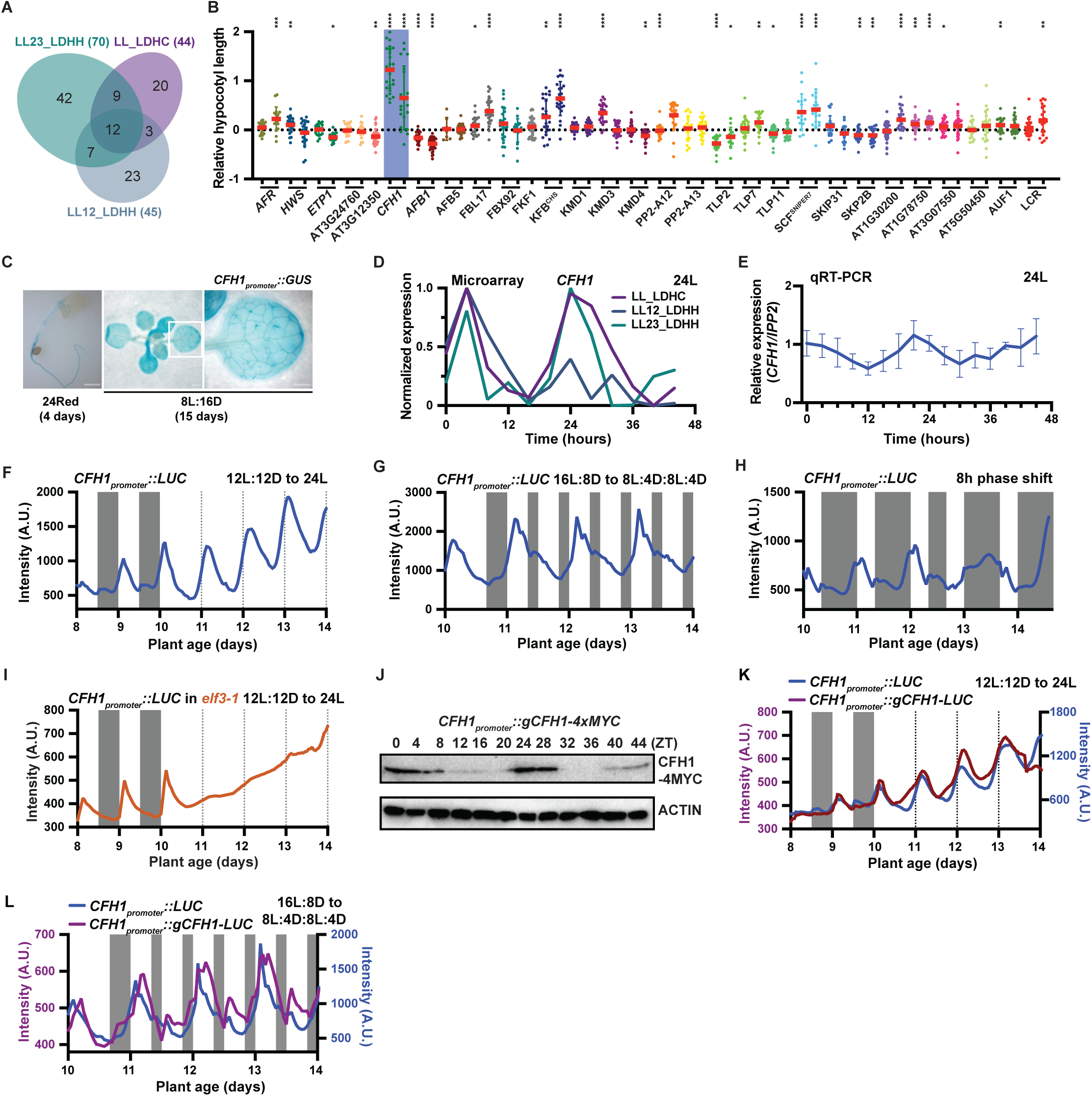
*CFH1* encodes a morning phased F-box protein that regulates hypocotyl length. (**A**). Venn diagram of 31 circadian clock-regulated F-box genes that show rhythmicity in at least two of the three microarray datasets in constant white light condition: LL_LDHC, LL12_LDHH, and LL23_LDHH (*6*). (**B**). Hypocotyl length of F-box decoy transgenics. Hypocotyl length is normalized to wild-type (Col). *CFH1* decoy lines highlighted in blue. Red line indicates average hypocotyl length. Error bars indicate SD (n = 7 – 50 seedlings). *, p ≤ 0.05; **, p ≤ 0.01; ***, p ≤ 0.001; ****, p ≤ 0.0001 (Welch’s t-test). (**C**). GUS staining of the *CFH1_promoter_::GUS* transgenic line. Left: 4-day-old seedling grown in constant red light. Scale bar = 0.5 mm. Middle, 15-day-old seedling grown in 8L:16D. Right: zoom-in view of the white box area in the middle image. Scale bar = 1 mm in the middle and right images. (**D**). Normalized expression pattern of *CFH1* in continuous white light from three microarray datasets. (**E**). qRT-PCR of *CFH1* expression from wild-type (Col) plants grown in constant white light. *IPP2* was used as an internal control. (**F-H**). Traces of *CFH1_promoter_::Luciferase* expression from plants grown in (**F**) 12L:12D to 24L, (**G**) 16L:8D to 8L:4D:8L:4D, and (**H**) 8L:16D with 8 h dawn phase shift at day 12. Gray shading represents dark periods. Lines represent intensity traces. (**I**). Traces of *CFH1_promoter_::Luciferase* in *elf3-1* mutant grown in 12L:12D and shifted to 24L. (**J**). Immunoblot detecting CFH1-4MYC driven by *CFH1* native promoter (*CFH1_promoter_::gCFH1-4MYC*). Two-day time course samples were collected from 9-day-old plants grown under constant white light. Actin was used as a loading control. (**K-L**). Traces of *CFH1_promoter_::gCFH1-Luciferase* and *CFH1_promoter_::Luciferase* from plants grown in (**K**) 12L:12D and shifted to 24L and (**L**) 16L:8D to 8L:4D:8L:4D.

The circadian clock regulates light sensing in plants, and we wanted to determine if any of these 31 clock regulated F-box genes are involved in photomorphogenesis. Studies of F-box genes can be hampered by genetic redundancy, but our lab developed a reverse genetic “decoy” strategy to overcome this problem (*11–15*). To create decoys, we express the protein recognition domains of E3 ubiquitin ligases without the F-box domain that recruits the ubiquitylation machinery, causing the decoy to stabilize, rather than degrade, their targets. We were able to create decoys for 29 of the 31 identified clock-regulated F-box genes, and we measured hypocotyl length in these lines in a 12 hour light: 12 hour dark (12L:12D) condition (Fig. 1B). *AT2G32560*, *F BOX-LIKE1* (*FBL17*), *KELCH DOMAIN-CONTAINING F-BOX PROTEIN* (*KFB^CHS^*), *SCF^SNIPER7^*, and *AT1G78750* decoy transgenic lines have hypocotyls longer than wild type, while *AUXIN SIGNALING F BOX PROTEIN 1* (*AFB1*), *TUBBY LIKE PROTEIN 2* (*TLP2*), and *ARABIDOPSIS HOMOLOG OF HOMOLOG OF HUMAN SKP2 2* (*SKP2B*) have hypocotyls shorter than wild type (*16*). Consistent with previous studies, *AFB1* is an auxin receptor that is known to control auxin sensitivity and hypocotyl growth (*17*). Notably, *AT2G32560* decoy transgenics exhibit the longest hypocotyl phenotypes. This gene has not been studied previously, and it has an uncharacterized C-terminal domain in addition to the F-box domain. Due to the strong rhythmic expression and large effects on hypocotyl length, we selected it for follow-up studies and named it *CLOCK-REGULATED F-BOX WITH A LONG HYPOCOTYL 1* (*CFH1*).

### *CFH1* is morning-phased

The *CFH1* decoy transgenics have hypocotyl defects but are expressed under a constitutive promoter. To ensure that CFH1 is expressed in the hypocotyl where we see the greatest defect, we determined the spatial expression pattern of *CFH1* using a transgenic line expressing β-glucuronidase under the control of *CFH1* promoter (*CFH1_promoter_::GUS*) (Fig. 1C). *CFH1* is expressed in the hypocotyl under red light at an early developmental stage (4 days) and also widely expressed in all tissues later in development (15 days).

Next, we wanted to further investigate the temporal expression of *CFH1.* Microarray data suggested that *CFH1* exhibits rhythmic expression in continuous light, phased at ZT 2-4 (probe 267116_at) (Fig. 1D). We tested and confirmed rhythmic expression of *CFH1* by qRT-PCR (Fig. 1E). To monitor the expression of *CFH1* more precisely, we generated a transgenic plant expressing the *Luciferase* gene under the control of the *CFH1* promoter (*CFH1_promoter_::LUC*) and measured luminescence from these plants in a circadian time course. We grew the plants in 12L: 12D then shifted them to constant light (12L:12D to 24L) (Fig. 1F). The pattern generated from this experiment was consistent with those seen in microarray and qRT-PCR. *CFH1* expression is rhythmic (RAE = 0.12) with a period of 23.88 hours and a phase at ZT 3.92 as calculated by the BioDare2 platform (biodare2.ed.ac.uk) (*18*). We next tested *CFH1* expression under 16L:8D, 12L:12D, 8L:16D, 16L:8D switched to 8L:16D, and 8L:16D switched to 16L:8D (Fig. S2A-B). *CFH1* expression consistently peaks in the early morning, showing that light dark cycles have little effect on the phasing of *CFH1*. qRT-PCR also confirmed the morning-phased expression pattern of *CFH1* in 12L:12D (Fig. S2C). We then grew the plants in a skeleton photoperiod of 8L:4D:8L:4D (Fig. 1G), and we found that the peak of *CFH1* remained at the first “dawn” of each 24 hour period. We then performed a phase shift experiment by growing the plants in 8L:16D and then advancing the phase of dawn by 8 hours on day 12 (Fig. 1H). Clock regulated genes require time to entrain to a new dawn after a phase shift. On day one after the phase shift, we observed a low amplitude and delayed phase of *CFH1* expression pattern. On day two after the phase shift, *CFH1* expression achieved its normal phase showing re-entrainment. The clock dampens quickly in continuous dark but can be partially restored by the addition of exogenous sucrose (*19*). We next tested *CFH1* expression in darkness and found that the rhythmicity of *CFH1* was dampened by the second day but that rhythmicity could be restored by adding sucrose (Fig. S2D). Together these results suggest that *CFH1* expression is regulated by the circadian clock but not diurnal cycles. To confirm that the canonical plant circadian clock regulates *CFH1* expression, we crossed the *CFH1_promoter_::LUC* to the arrhythmic clock mutant *elf3-1* (Fig. 1I). The *elf3-1* mutant caused arrhythmicity of *CFH1*, confirming that it is regulated by the circadian clock.

We next tested whether CFH1 protein levels oscillate. We generated transgenic plants expressing the CFH1 protein fused with luciferase or MYC expressed under the control of the *CFH1* promoter (*CFH1_promoter_::gCFH1-LUC* and *CFH1_promoter_::gCFH1-4MYC*), and monitored protein luminescence or abundance in continuous light (Fig. 1J-K). Similar to *CFH1* mRNA, CFH1 protein has an RAE of 0.16 and peaks in the early morning, phased at ZT 4.03 with a period of 25.01 hours. We also grew the *CFH1_promoter_::gCFH1-LUC* reporter in the skeleton photoperiod of 8L:4D:8L:4D (Fig. 1L), and the protein maintained its phase, showing that the protein is not influenced by diurnal cycles. Taken together, our results demonstrate that both *CFH1* mRNA and protein levels are regulated by the circadian clock and phased to the early part of the day.

### CFH1 regulates red light morphogenesis

Light fluence and duration affect photomorphogenic hypocotyl elongation. We tested whether the *CFH1* decoy had photoperiod- or fluence-specific effects on photomorphogenesis. We assessed hypocotyl length under a range of white light photoperiods and constant blue or red light (Fig. 2A). The *CFH1* decoy transgenics have long hypocotyls in 8L:16D, 12L:12D, 16L:8D, and 24L and small but significant changes in hypocotyl length in 24Blue and 24D. Strikingly, the largest difference in hypocotyl length was observed in constant red light (1.6-fold greater than wild type). Next, we identified two *cfh1* insertion mutants *cfh1-1* (CS853743) and *cfh1-2* (CS429829) and generated a CRISPR deletion mutant (*cfh1c*) (Fig. S3A). We confirmed that the expression of *CFH1* was compromised in the mutants (Fig. S3B-C) and then used the *cfh1-1* mutant to test hypocotyl length in the same conditions as the decoy transgenics (Fig. 2B). We found that *cfh1-1* mutant exhibited similar hypocotyl defects as the *CFH1* decoy transgenic plant, confirming its role in red light photomorphogenesis. Additionally, *cfh1-1* can be complemented by expressing full-length *CFH1* driven by the native promoter (Fig. 2C), and *cfh1-2* and *cfh1c* mutant plants also have long hypocotyls in red light (Fig. S3D). Together our results show that *CFH1* regulates hypocotyl elongation particularly in red light.

**Fig. 2.**
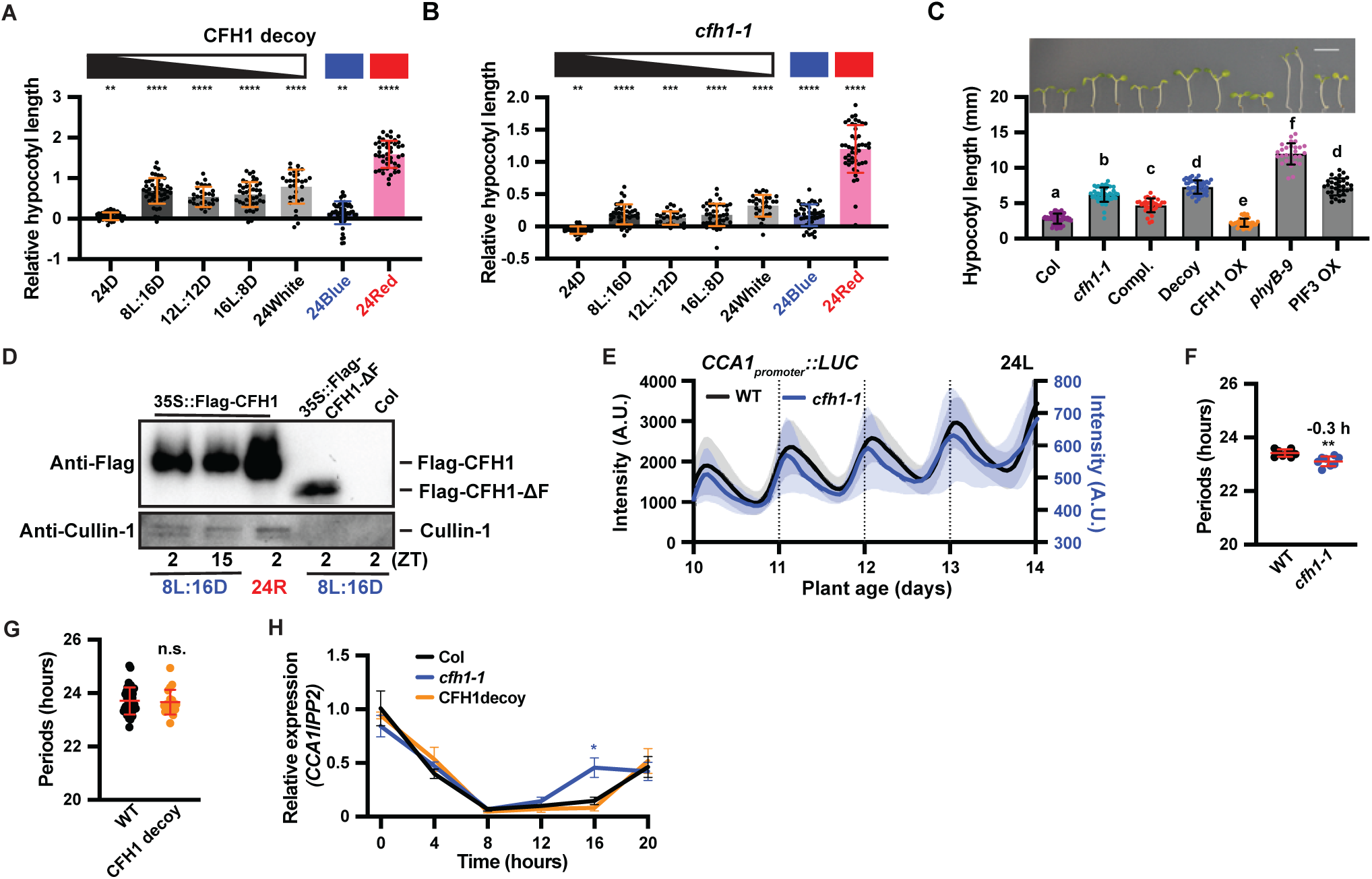
CFH1 is part of an SCF complex that regulates hypocotyl length. (**A-B**). Hypocotyl length of (**A**) *CFH1* decoy and (**B**) *cfh1-1* mutant in 24D, 8L:16D, 12L:12D, 16L:8D, 24L, 24Blue, and 24Red. The hypocotyl length was normalized to wild-type (Col). Error bars indicate SD (n = 29 – 51 seedlings). **, p ≤ 0.01; ***, p ≤ 0.001; ****, p ≤ 0.0001 (Welch’s t-test). (**C**). Hypocotyl length and representative images of 4-day-old wild-type (Col), *cfh1-1* mutant, *cfh1-1* complementation line (Compl.), *CFH1* decoy (Decoy), and *CFH1* OX grown in 24Red. *phyB-9* and *PIF3* OX plants were grown as controls. Different letters indicate statistically significant differences as determined by one-way ANOVA followed by Dunnett’s T3 multiple comparison test. P ≤ 0.05. Error bars indicate SD (n = 20 – 63 seedlings). Scale bar = 5 mm. (**D**). Co-IP showing interaction between CFH1 and the SCF scaffold protein Cullin-1. Immunoprecipitation was performed with total proteins extracted from 12-day-old wild-type (Col), *35S::Flag-CFH1,* or *35S::Flag-CFH1ΔF* plants grown in 8L:16D or 24Red. Samples were collected at ZT2 or ZT15 as indicated. (**E**). *CCA1_promoter_:: Luciferase* traces from wild type (Col) or *cfh1-1* mutant plants grown in constant white light. Shading indicates SD. (**F**). Clock period of *CCA1_promoter_:: Luciferase* in wild-type (Col) and *cfh1-1* mutant plants calculated from E. Number on the data indicates hours of period change. Error bars indicate SD (n = 8 – 10 plants). **, p ≤ 0.01 (Welch’s t-test). (**G**). Clock period of *CCA1_promoter_:: Luciferase* in wild-type (Col) and *CFH1* decoy. Error bars indicate SD (n = 19 – 40 plants). n.s., not statistically significant (Welch’s t-test). (**H**). qRT-PCR of *CCA1* in wild-type (Col), *cfh1-1* mutant, and *CFH1* decoy plants grown in constant white light. *IPP2* was used as an internal control. Error bars indicate SD (n = 3 replicates). *, p ≤ 0.05 (Welch’s t-test). Star color indicates the statistical analysis between the colored genotype (blue for *cfh1-1*; orange for *CFH1* decoy) and wild-type (Col).

We next generated a *CFH1* overexpression (*CFH1* OX) transgenic line by expressing full length *CFH1* under the *35S* promoter (Fig. 2C). These plants have shorter hypocotyls in red light than the wild-type plants, in contrast to the *CFH1* decoy transgenics that have longer hypocotyls in red light. This indicates that the F-box domain is important for CFH1 function. To confirm the role of the F-box domain in SCF formation, we performed co-immunoprecipitations (co-IP) using an affinity-tagged CFH1 over-expression line (*35S::Flag-CFH1*) and a *CFH1* decoy line which also contains an affinity tag (*35S::Flag-CFH1ΔF*) (Fig. 2D). We immunoprecipitated the full length and F-box deleted CFH1 proteins and used western blotting to detect the SCF scaffold protein Cullin-1. Cullin-1 was detected from the full length CFH1 IP samples in all timepoints but was not detected in the *CFH1* decoy transgenic because the F-box domain is removed. Our results show that proper *CFH1* expression and formation of an SCF complex are required for its role in hypocotyl growth regulation.

### CFH1 is a circadian clock output regulator

Circadian clock mutants can have red light hypocotyl growth defects, similar to the *cfh1-1* mutant. Thus, we wanted to test if *CFH1* is directly regulating the circadian clock, part of a clock feedback loop, or is a *bona fide* output regulator. First, we crossed the *cfh1-1* mutant, or transformed *CFH1* decoy, to a transgenic line expressing *Luciferase* under the *CCA1* promoter (*CCA1_promoter_::LUC*) (*20, 21*). We then grew the plants in constant light and measured the period (Fig. 2E-G). The period of *CCA1* in the *cfh1-1* mutant was slightly shorter than wild type (−0.3 h), but the same effect was not seen in the *CFH1* decoy transgenic lines. We next performed qRT-PCR to measure the expression level of *CCA1* (Fig. 2H), *LHY*, *PRR9*, *PRR7*, *PRR5*, *TOC1*, and *GI* (Fig. S4) in the *cfh1-1* mutant and *CFH1* decoy transgenics in constant light. *CCA1* and *LHY* have a small increase in expression at ZT16 only in the *cfh1-1* mutant, and *TOC1* has higher amplitude only in the *CFH1* decoy transgenic. We next tested whether CFH1 interacts with core clock components using yeast two-hybrid (Fig. S5A). We found no interactions between CFH1 and the core clock machinery. Our results show a lack of consistency between the *cfh1-1* and *CFH1* decoy transgenic in clock gene expression and a lack of interaction with clock components. Thus, we conclude that CFH1 is likely a *bona fide* output regulator rather than directly or indirectly controlling clock function.

### CFH1 controls hypocotyl growth through degradation of PIF3

We next wanted to determine how CFH1 controls hypocotyl elongation. The phenotypic effects of CFH1 in the regulation of hypocotyl length in red light are reminiscent of the phyB-PIF signaling model (*22, 23*) (Fig. 2A-C). Therefore, we used yeast two-hybrid to test if CFH1 interacts with Hy5 or PIFs (PIF1, PIF3, PIF4, and PIF5), red light transcription factors that regulate hypocotyl length (Fig. 3A). We found that the *CFH1* decoy was able to specifically interact with PIF3, but not other PIFs or Hy5. Interestingly, full length CFH1 did not interact with PIF3 (Fig. S5B), suggesting that CFH1 may form an SCF complex and degrade PIF3 in yeast. We next tested the CFH1-PIF3 interaction *in vivo* using split luciferase and co-IP (Fig. 3B-C). In tobacco leaves, luminescence was detected when PIF3-nLUC and cLUC-CFH1ΔF were co-expressed but not when PIF5-nLUC and cLUC-CFH1ΔF were co-expressed, confirming that CFH1 and PIF3 interact specifically (Fig. 3B). co-IP again confirmed the *in planta* interaction between CFH1 and PIF3 (Fig. 3C).

**Fig. 3.**
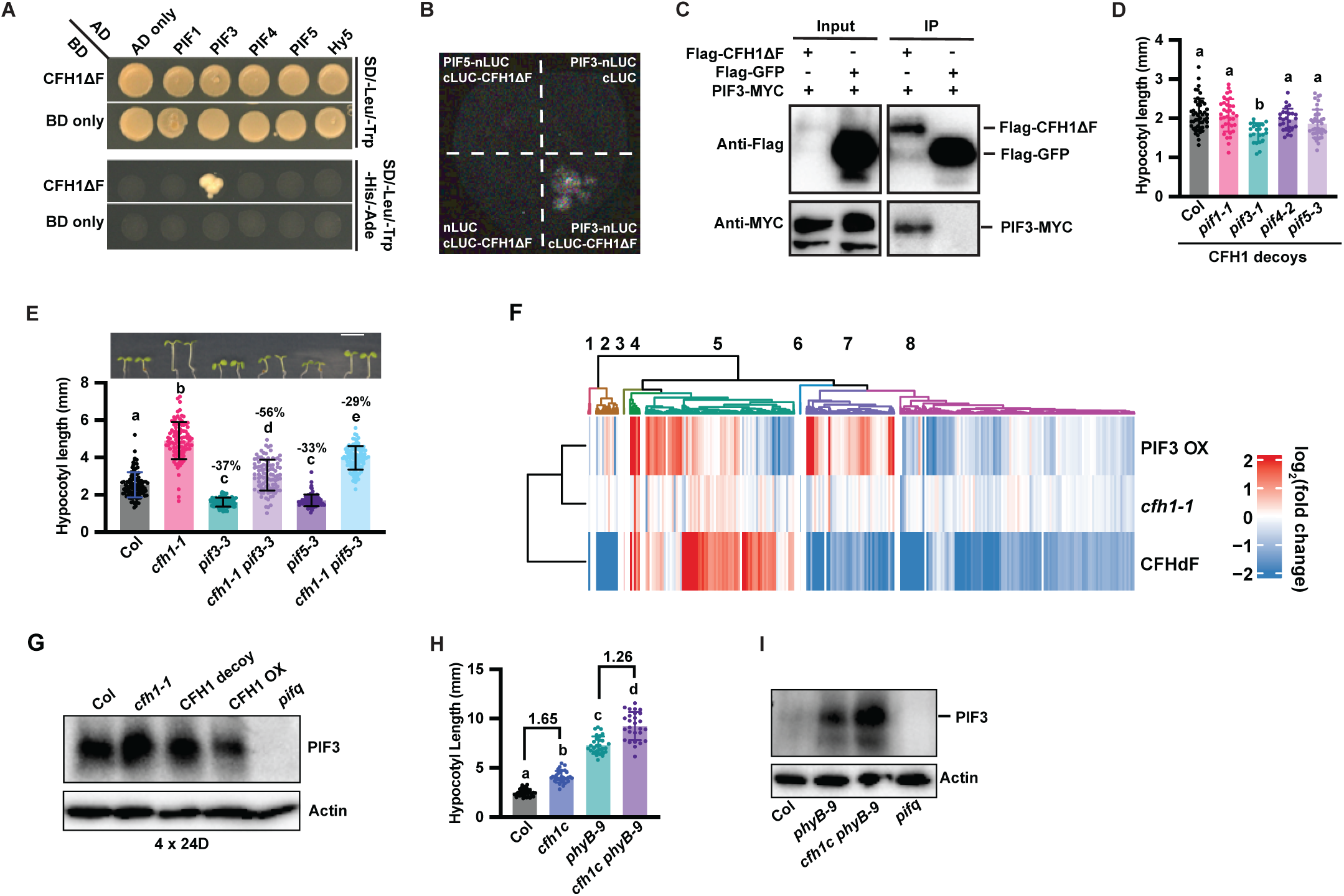
CFH1 works in parallel to PhyB to regulate PIF3 stability. (**A**). Yeast two-hybrid assay. Yeast was transformed with GAL4 DNA-binding domain (BD) fused to CFH1ΔF (BD-CFH1ΔF) and GAL4 activation domain (AD) fused to PIF1, PIF3, PIF4, PIF5, or Hy5 and grew in SD-Leu-Trp medium for autotrophic selection and on SD-Leu-Trp-His-Ade medium to test for interactions. (**B**). Split Luciferase assay. Constructs of PIF3 fused to nLUC and CFH1ΔF fused to cLUC were co-expressed in tobacco leaves. PIF5 fused to nLUC or empty vector were used as negative controls. (**C**). Co-IP assay. Total proteins were extracted from 4-day-old dark grown plants of *35S::PIF3-MYC* crossed to *35S::Flag-CFH1ΔF* or *35S::Flag-GFP* plants. (**D**). Hypocotyl length of 7-day-old T1 *CFH1* decoy plants in wild-type (Col), *pif1-1*, *pif3-1*, *pif4-2*, and *pif5-3* mutant background. Error bars indicate SD (n = 23 – 47 seedlings). (**E**). Hypocotyl length and representative images of 4-day-old Col, *cfh1-1*, *pif3-3*, *cfh1-1 pif3-3*, *pif5-3*, and *cfh1-1 pif5-3* plants grown in 24Red. Percent growth inhibition is indicated for *pif3-3* and *pif5-3* relative to the wild-type (Col) as well as *cfh1-1 pif3-3* and *cfh1-1 pif5-3* relative to parents were indicated on the column. Error bars indicate SD (n = 98 – 100 seedlings). Scale bar = 5 mm. (**F**). RNA-seq heatmap depicting magnitude of log_2_(fold change) of differentially expressed genes between *cfh1-1*, *CFH1* decoy, and PIF3 OX lines against the wild-type (Col). Hierarchical clustering is presented as dendrograms on the top and the left. (**G**). Immunoblot detecting endogenous PIF3 protein levels in *cfh1-1* mutant, *CFH1* decoy, and *CFH1* OX plants grown in constant dark for 4 days. Wild-type (Col) and *pifq* mutant were used as controls. Actin was used as loading control. (**H**). Hypocotyl length of wild-type (Col), *cfh1c*, *phyB-9*, and *cfh1c phyB-9* mutants plants grown in 24Red for 4 days. Error bars indicate SD (n = 27 – 35 seedlings). Numbers above the data indicate fold change. In D, E, and H, different letters indicate statistically significant differences as determined by one-way ANOVA followed by Dunnett’s T3 multiple comparison test. P ≤ 0.05. (**I**). Immunoblot detecting endogenous PIF3 protein levels in wild-type (Col), *phyB-9* mutant, *cfh1c phyB-9,* and *pifq* mutant plants grown in 8Red:16D for 4 days. On the 4^th^ day, plants were given an extended night and samples were collected at ZT 1 in the dark.

We next wanted to test this interaction genetically. First, we generated *CFH1* decoy transgenic lines in wild type (Col), *pif1-1*, *pif3-1*, *pif4-2*, and *pif5-3* mutants backgrounds and compared the hypocotyl lengths of T1 transgenic lines grown in 12L:12D (Fig. 3D). Expressing the *CFH1* decoy was able to cause hypocotyl lengthening in wild type and all mutant backgrounds except *pif3-1*. Next, we crossed the *cfh1-1* mutant to *pif3-1* and *pif5-3* and measured the hypocotyls of plants grown in red light (Fig. 3E). *pif3-3* and *pif5-3* single mutants cause similar hypocotyl shortening in red light (−37% and −33%). In the *cfh1* mutant background, *pif3-1* causes greater hypocotyl shortening (−56%) than in the wild-type background, suggesting a non-additive genetic interaction between the two genes, while *pif5-3* causes a similar percent decrease (−29%) in hypocotyl length between wild type and the *cfh1-1* mutant, demonstrating an additive non-genetic interaction. These data support the idea that CFH1 is specifically regulating PIF3 to control hypocotyl length.

The physical and genetic interactions between CFH1 and PIF3 suggest that CFH1 should regulate a similar subset of genes as PIF3. To test this idea, we performed RNA-seq at ZT1 in constant red light, a time point close to the *CFH1* expression peak. We collected RNA samples from wild type, the *chf1-1* mutant, the *CFH1* decoy transgenic line, and a *PIF3* over-expression (*PIF3* OX) transgenic line (Fig. S6A-E, Table S2). We found 17 down- and 2 up-regulated genes in the *cfh1-1* mutant; 1992 down- and 636 up-regulated genes in the *CFH1* decoy transgenic; and 452 down- and 462 up-regulated genes in the *PIF3* OX transgenic. We next performed clustering of the differentially expressed genes from the three genotypes (Fig. 3F). While the magnitude of expression changes varied between the three genotypes, many genes were co-repressed (cluster 8: 1466 genes) and co-induced (cluster 4: 64 genes) between the three genotypes (Fig. 3F, S6D-E, Table S2). This provides further evidence that CFH1 is regulating PIF3 function.

CFH1 and PIF3 interact physically, and CFH1 can form an SCF complex. We next wanted to determine if PIF3 protein stability is regulated by CFH1. We acquired an antibody to the native PIF3 protein and performed western blots on wild-type, *cfh1-1* mutant, *CFH1* decoy transgenic, *CFH1* OX, and the *pifq* mutant (*24*) (Fig. 3G). We grew the plants in 4 days of constant dark to induce accumulation of PIF3 and found the *cfh1-1* mutant and the *CFH1* decoy transgenic line had increased PIF3 protein levels. Conversely, the *CFH1* OX transgenic line had lower PIF3 levels. These results suggest that CFH1 regulates the degradation of PIF3.

Two E3 ligase families have been shown to regulate PIF3 degradation in response to red light (*25, 26*). We next wanted to test whether the circadian clock and CFH1 work in parallel to red light signaling to regulate PIF3 protein stability. To test this, we generated a CRISPR deletion mutation in *CFH1* in the *phyB-9* mutant background. We measured hypocotyl length and found that the *phyB-9 cfh1c* double mutant had longer hypocotyls than *phyB-9* alone (Fig. 3H). Furthermore, we measured PIF3 protein abundance and found that PIF3 accumulates to a greater extent in the *phyB-9 cfh1c* double mutant than in the single *phyB-9* mutant (Fig. 3I). These results show PIF3 protein stability is a convergence point for red light signaling and the circadian clock to control photomorphogenesis.

## Discussion

Eukaryotic circadian clocks are composed of transcriptional feedback loops that are required for rhythmicity of the clock itself but also impose rhythmicity on output processes (*21, 27*). In the last decades, it has become increasingly clear that the transcription factors that make up the core of the circadian clock require extensive post-translational modification and targeted degradation by the ubiquitin proteasome system to maintain 24-hour rhythms (*1, 3*). What has been less clear is the extent to which the ubiquitin proteasome system is recruited by the circadian clock to regulate output processes. Here we isolated circadian clock regulated E3 ubiquitin ligases and discovered a degradation mechanism that tethers the circadian clock to photomorphogenesis.

There are multiple nodes of connection between the circadian clock and red light signaling, highlighting the importance of proper timing of light responses throughout the day. For instance, the evening complex (EC) associates with phytochromes to occupy target gene promoters and control gene expression (*28*). The core clock components PRR7 and PRR5 directly bind PIFs to repress their transcriptional activity (*29*), and TOC1 interacts with PIF4 to medicate the circadian gating of thermo-responsive growth (*30*). The connection reported here is unique. There is little or no clock control of *PIF3* mRNA, rather the clock connects to PIF3 through regulated protein degradation (*31*). PIF3 in turn regulates photosynthesis and growth, and unlike other PIFs, does not feed back into the clock (*32–34*). This finding demonstrates that E3 ubiquitin ligases can act as a tethering mechanism between the clock and *bona fide* outputs.

Two additional degradation mechanisms regulate PIF3 stability, but both are controlled by red light signaling through phytochrome (*25, 26, 35*). Here we show that the clock and CFH1 function in parallel to red light signaling to regulate PIF3 stability and photomorphogenesis (Fig.3I-J; Fig. S7). The importance of the clock lies in its ability to retain its daily rhythms when environmental signals fluctuate. Thus, we propose that the CFH1 tether to PIF3 “guarantees” the plant a morning state even when phytochrome activity is compromised.

Interestingly, our genetics experiment suggests that PIF3 may not be the only target of CFH1 (Fig. 3E), and it will be important in the future to identify the full range of CFH1 targets. This can be facilitated by the decoy approach which allows for immunoprecipitation followed by mass spectrometry to identify protein interacting partners (*11–15, 36*). Additionally, CFH1 has two homologs in Arabidopsis and orthologs in other species, indicating that plants have retained this gene throughout evolution and highlighting its importance.

Here we characterize one post-translational tether between the circadian clock and an important output in plants. This work opens the possibility that there are many additional protein degradation-based clock tethering mechanisms that exist and provides a roadmap for identifying them. This also highlights that the connections between the clock and outputs are complicated hierarchical networks that are underexplored.

## ACKNOWLEDGEMENTS

We would like to thank Dr. Dmitri Nusinow and Dr. Jie Dong for clock- and PIFs-related mutants as well as transgenic plants, Dr. Ning Wei for Cullin-1 antibody, Dr. Geoffrey Thomson and Dr. Valentin Joly for CRISPR-related technical support, and Daniel Tartè for luciferase imaging support. We would also like to thank Sandra Pariseau and Jenny Pengsavath for administrative support. Additionally, we would like to thank Chris Bolick, Nathan Guzzo, and the staff at Marsh Botanical Gardens for their support in maintaining plant growth spaces. We would also like to thank Dr. Man-Wah Li, Dr. Qingqing Wang, Morgan Vanderwall, Anxu Xu for insightful discussions and critical reading of the manuscript. This work was supported by the National Institutes of Health (R35 GM128670) to J.M.G.. W.L. was supported by the Forest BH and Elizabeth DW Brown Fund Fellowship.

## AUTHOR CONTRIBUTIONS

W.L., H.L., C.C.L., C.A., J.D., J.H, M.C., and J.M.G. designed the experiments. W.L., H.L., C.C.L., C.A., J.D., and J.H performed the experiments and experimental analyses. W.L. C.C.L., and J.M.G. wrote the article.

## Materials and methods

### Arabidopsis materials

The *Arabidopsis* seeds of Col-0, *cfh1-1* (CS853743), *cfh1-2* (CS429829) were obtained from ABRC. The *elf3-1* mutant seed was obtained from Dr. Dmitri Nusinow (*38*). *phyB-9*, *pif3-1*, *pif3-3*, *pif4-2*, *pif5-3*, *pifq*, and *PIF3-MYC* over-expression (*PIF3* OX) transgenic lines were obtained from Dr. Jie Dong (*25, 39*). *CCA1_promoter_::luciferase* was described previously (*20*). 29 F-box decoy transgenic lines, *CFH1_promoter_::luciferase*, *CFH1_promoter_::gCFH1-luciferase*, *CFH1_promoter_::gCFH1-MYC*, *CFH1_promoter_::gCFH1-*GFP in Col or *cfh1-*1 mutant background, *35S::Flag-CFH1* (*CFH1* OX), *CFH1 crispr* (*cfh1c*) in Col and *phyB-*9 mutant background were generated in this study. The *CFH1_promoter_::luciferase* in *elf3-1* mutant, *CCA1_promoter_::luciferase* in *cfh1-1* mutant, *cfh1-1 pif3-3*, and *cfh1-1 pif5-3* double mutants were generated by crossing and homozygous lines were confirmed by phenotypes, luciferase imaging and genotyping. The primers used for genotyping are listed in table S3.

### Arabidopsis growth condition

Arabidopsis seeds were surface sterilized for 20 min in 70% ethanol with 0.1% Triton X-100 then sown on freshly poured ½ MS plates, pH 5.7, (Cassion Laboratories, cat. # MSP01) and 0.8% bacteriological agar (AmericanBio cat. # AB01185) without sucrose. For hypocotyl assays, the seeds were stratified in the dark for two days at 4 °C and then transferred into 22 °C, 70 µmol/m^2^/s constant red light for 4 days or 150 µmol/m^2^/s white light with various photoperiods as indicated for 7 days, to measure hypocotyl length. For luciferase imaging, after stratification, seeds were transferred into 22 °C, 12L:12D illuminated by 150 µmol/m^2^/s white light for seven days and then transferred to various photoperiods for given experiments as indicated for imaging. For RNA-sequencing, after germinated for seven days in 12L:12D (150 µmol/m^2^/s), seedlings were transferred to 70 µmol/m^2^/s constant red light. Samples were collected at day 12 ZT1.

### Plasmid construction

For F-box genes over-expression or decoy constructs, the coding sequence of the F-box gene with or without F-box domain were obtained by PCR using Col cDNA as template, inserted into pENTR/D-TOPO (Invitrogen, cat. # K240020) and then transferred into PB7-HFN destination vectors using LR recombination (*40*). To generate the *CFH1_promoter_::LUC* and *CFH1_promoter_::GUS* construct, a 1980 bp promoter sequence upstream of the *CFH1* coding sequence, including 5’ UTR, was obtained by PCR and inserted into pENTR/D-TOPO vector and then transferred into the pFLASH and pMDC164 destination vectors to drive the luciferase (*21*) and the GUS (*41*), respectively. To generate the *CFH1_promoter_::gCFH1* tag constructs, the *CFH1_promoter_::gCFH1* fragment was generated from PCR using Col genomic DNA as the template, inserted into pENTR/D-TOPO and then transferred into pGWB4, pGWB16, and pGWB435 destination vectors using LR recombination to generate *CFH1_promoter_::gCFH1-GFP, CFH1_promoter_::gCFH1-4MYC,* and *CFH1_promoter_::gCFH1-LUC*, respectively (*42*). The *cfh1c* deletion mutant was generated by CRISPR using two guide RNA in *CFH1* as described previously (*43*). For split luciferase assay, PIF3 or PIF5 was subcloned into pGWB-nLUC (Addgene #174050) and CFH1ΔF was inserted into pGWB-cLUC (Addgene #174051) (*44*). The primers used for cloning are listed in table S3.

### Luciferase imaging

Luciferase imaging and data analysis were performed as previously described (*7*). Briefly, seven-day old seedlings grown in 12L:12D at 22 °C for 7 days were transferred onto a 10 x 10 grid freshly poured 100 mm square ½ MS plates with or without added sugars as indicated for given experiments. Seedlings were then treated with 5 mM D-luciferin (Cayman Chemical Company, cat. # 115144-35-9) dissolved in 0.01% TritonX-100, and imaged at 22 °C under the indicated conditions for 7 days.

### qRT-PCR

For qRT-PCR experiments, total RNA was extracted from Arabidopsis seedlings grown in indicated conditions with RNeasy Plant Mini Kit (QIAGEN cat. # 74904) and then treated with DNase (QIAGEN, cat. # 79254). The subsequent reverse-transcription and conditions for qRT-PCR reactions were described previously with minor modifications (*11*). Briefly, four hundred nanograms of total RNA were used for reverse-transcription using iScript™ Reverse Transcription Supermix for RT-qPCR (Bio-Rad, cat. # 1708841). iTaq Universal SYBR Green Supermix was used for qRT-PCR reaction (Bio-Rad, cat. # 1725121). *IPP2* (AT3G02780) or *UBQ10* (AT4G05320) was used as internal controls as indicated. The relative expression represents means of 2^(−ΔCT)^ from three biological replicates, in which ΔCT = (CT of Gene of Interest – CT of internal control). The primers used for qRT-PCR are listed in table S3.

### GUS histochemical analysis

For GUS assay, the *CFH1_promoter_::GUS* transgenic plant was grown in 70 µmol/m^2^/s constant red light for 4 days or 150 µmol/m^2^/s white light with photoperiod of 12L:12D for 12 days and then transferred to 8L:16D for 3 more days. The plant was freshly harvested and stained at 37 °C over night with 2 mM 5-bromo-4-chloro-3-indolyl-beta-D-glucuronic acid (X-gluc) in 100 mM potassium phosphate buffer, pH 7.0, containing 0.1% (v/v) Triton X-100, 1 mM K_3_Fe(CN)_6_ and 10 mM EDTA. Tissues were cleared before observation by washing with 70% and 50% (v/v) ethanol.

### Yeast two-hybrid

The yeast two-hybrid assay was performed on synthetic dropout medium as described previously (*11*). Briefly, indicated proteins were fused to the GAL4-BD in pGBKT7-GW vectors or GAL4-AD in pGADT7-GW vectors by GATEWAY cloning. The interactions were tested on synthetic dropout medium as indicated.

### Co-IP

For CFH1-Cullin-1 co-IP, wild-type (Col), *35S::Flag-CFH1*, and *35S::Flag-CFH1ΔF* plants were grown in 12L:12D for 7 days and then transferred to constant red (70 µmol/m^2^/s) or 8L:16D (150 µmol/m^2^/s) for 4 days. Samples were collected at ZT2 or ZT15 at day 12. For CFH1-PIF3 co-IP, F2 population of *35S::Flag-CFH1ΔF* x *35S::PIF3-MYC* and *35S::Flag-GFPF* x *35S::PIF3-MYC* seeds were germinated in constant dark for 4 days. For both experiments, total proteins were extracted with SII buffer (100 mM sodium phosphate, pH 8.0, 150 mM NaCl, 5 mM EDTA, and 0.1% [v/v] Triton X-100) with cOmplete EDTA-free Protease Inhibitor Cocktail (Roche, catalog no. 11873580001), 1 mM PMSF, a PhosSTOP tablet (Roche, catalog no. 04906845001), and 50 µM MG132. The anti-FLAG antibodies (Millipore-Sigma F1804) were cross-linked to Dynabeads M-270 Epoxy (Thermo Fisher Scientific, catalog no. 14311D) for immunoprecipitation. Immunoprecipitation was performed by incubation with beads at 4°C for 2 h on a tube rocker and then washed three times with SII buffer.

### Immunoblotting

For the immunoblot assay in figure 3G and 3I, total proteins were extracted at indicated time as previously described (*45*), with extraction buffer consisted of 100 mM Tris-HCl, pH 7.5, 100 mM NaCl, 5 mM EDTA, pH 8.0; 5% SDS, 20% glycerol, 20 mM DTT, 40 mM β-mercaptoethanol, cOmplete EDTA-free Protease Inhibitor Cocktail (Roche), 2 mM PMSF, 80 μM MG132 (Sigma-Aldrich), a PhosSTOP tablet. The Flag-tagged proteins were then detected with Flag-HRP antibody (Sigma A8592, 1:5000); Endogenous PIF3 was detected with PIF3 antibody (Agrisera AS163954, 1:1000). Actin protein was detected with anti-Actin antibody (Millipore-Sigma MAB1501, 1:3000). For other immunoblotting, PIF3-MYC was detected with polyclonal MYC antibody (Sigma C3956, 1:3000); CFH1-MYC was detected with monoclonal MYC antibody 9E10 (Invitrogen MA1-980, 1:3000); Cullin-1 was detected with Cullin-1 antibody (*46*) (1:3000).

### RNA extraction and library preparation

RNA extraction and library preparation was performed as described previously with minor modifications (*47*). Briefly, the total RNA was extracted from approximately 200 mg of ground Arabidopsis seedlings using TRIzol reagent (ThermoFisher, 15596026) according to manufacturer’s protocol. RNA samples were treated with RNase-free DNase (QIAGEN, 79254) to remove DNA contaminants and further cleaned with RNA Clean & Concentrator-25 (ZYMO research #R1017). RNA-sequencing libraries were prepared and sequenced at the Yale Center for Genome Analysis. Samples were checked for quality with Agilent Bioanalyzer and those with > 7.0 RNA integration number were processed with the mRNA Seq Kit (Illumina, cat. # 1004814). 7 microliters of oligo dT on Sera-magnetic beads and 50 μL of binding buffer were used for isolation of mRNA, which was subsequently fragmented in the presence of divalent cations at 94^0^C. SuperScriptII reverse transcriptase (ThermoFisher, cat. # 18064014) was used for reverse transcription. The adaptor-ligated DNA was then amplified and purified with the Qiagen PCR purification kit (QIAGEN, 28104). Sequencing was performed the Illumina NovaSeq6000 platform with S1 flow cells in paired end mode at 100 base pairs.

### RNA-sequencing analysis

Trimmomatic (v.0.39) was used to trim sequencing adapters (ILLUMINACLIP:TruSeq3-PE-2.fa:2:30:10:1 LEADING:5 TRAILING:5 SLIDINGWINDOW:4:20 MINLEN:36) (*48*). The cleaned reads were aligned to the transcriptome (Ensembl version 57) with salmon (v.1.4.0) (*49*) using genomic sequence as decoy. Salmon-quantified transcripts were merged into genes with tximport (v1.26.1) and differential expression analysis was performed with DESeq2 (v1.38.3) (*50*). Only genes with at least 20 reads in at least 6 libraries were kept for analysis. Differential expression was defined at |log_2_(fold change)| > 1 and adjusted-*p*value < 0.05 (Benjamini Hochberg correction) using the Wald’s test. To identify potential CFH1-regulated genes downstream of PIF3, we identified PIF3-bound genes that were co-induced or co-repressed in the *PIF3 OX* transgenic line, the CFH1 decoy and the *cfh1-1* mutant, and differentially expressed in the *PIF3 OX* transgenic line and the CFH1 decoy. The PIF3-bound genes were reported from a ChIP-seq experiment performed and re-analyzed previously (*37, 51*). For gene expression clustering, the unshrunk log_2_(fold change) of gene expression against the wild type was used. Clustering was performed with the complete method and Euclidean distance using the hclust function in R. Heatmap was generated with ComplexHeatmap (*52*).

---

**Fig. S1.**
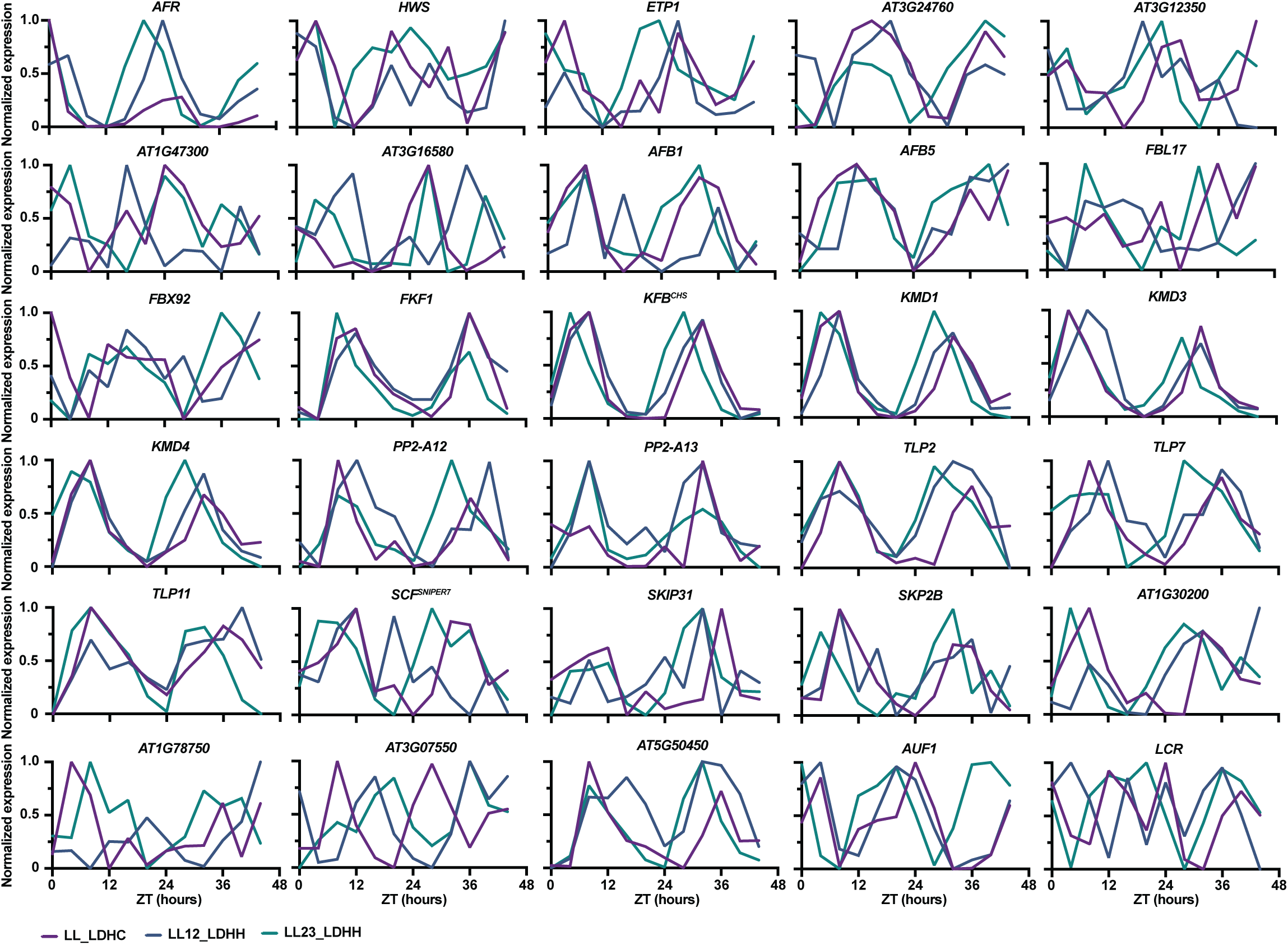
Normalized expression pattern of 30 circadian clock-regulated F-box genes in constant light conditions from three microarray datasets (LL_LDHC, LL12_LDHH, LL23_LDHH).

**Fig. S2.**
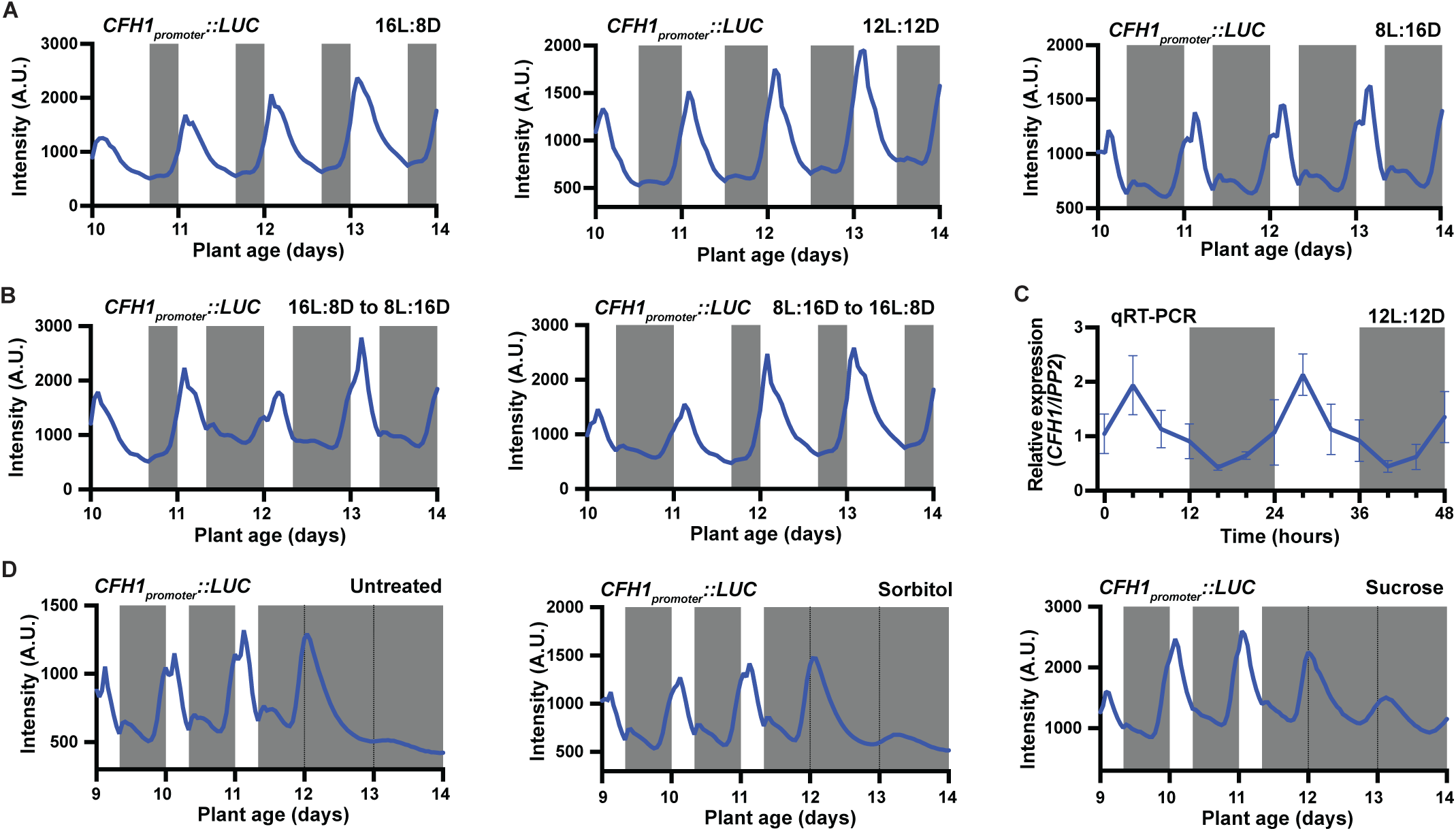
Expression pattern of *CFH1* in diurnal and constant dark conditions. (**A-B**). Traces of *CFH1_promoter_::Luciferase* from plants grown in (**A**) 16L:8D, 12L:12D, 8L:16D and (**B**) 16L:8D switched to 8L:16D and 8L:16D switched to 16L:8D. (**C**). qRT-PCR of *CFH1* expression in 12L:12D. (**D**). Traces of *CFH1_promoter_::Luciferase* from plants grown in 8L:16D and transferred to constant dark at day 12 with or without 90 mM sucrose or sorbitol as indicated.

**Fig. S3.**
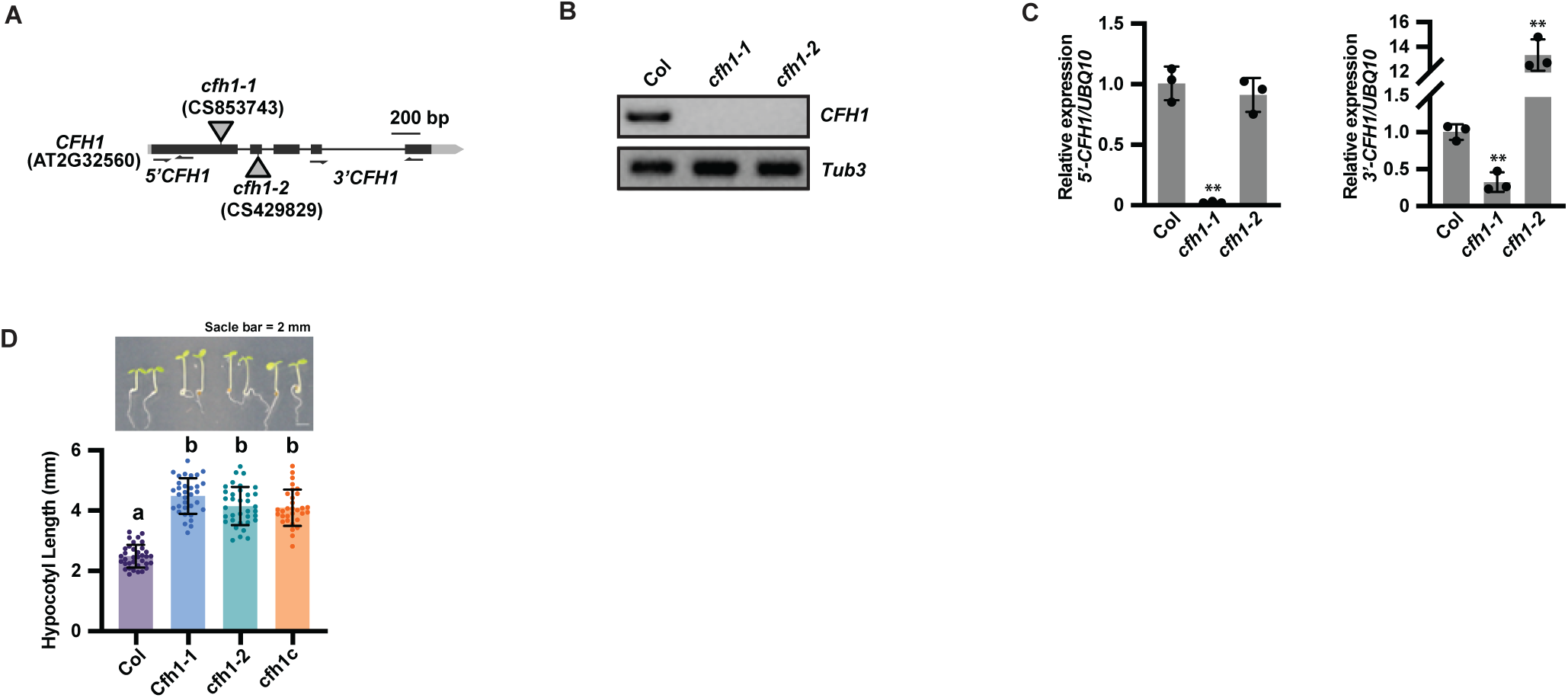
Characterization of *cfh1* mutants. (**A**). Schematic of *cfh1-1* and *cfh1-2* T-DNA insertion mutants. (**B**). Semi-quantitative PCR showing *CFH1* full length transcript was undetected *in cfh1-1* and *cfh1-2* mutants. *Tub3* was used as a control. (**C**). qRT-PCR of the 5’ and 3’ ends of *CFH1* transcripts in *cfh1-1* and *cfh1-2* mutants. *UBQ10* was used as a control. Error bars indicate SD (n = 3 replicates). **, p ≤ 0.01 (Welch’s t-test). (**D**). Hypocotyl length and representative images of wild-type (Col), *cfh1-1*, *cfh1-2*, and *cfhc* plants grown in 4 days of 24Red. Different letters indicate statistically significant differences as determined by one-way ANOVA followed by Dunnett’s T3 multiple comparison test. P ≤ 0.05. Error bars indicate SD (n = 31 – 35 seedlings). Scale bar = 2 mm.

**Fig. S4.**
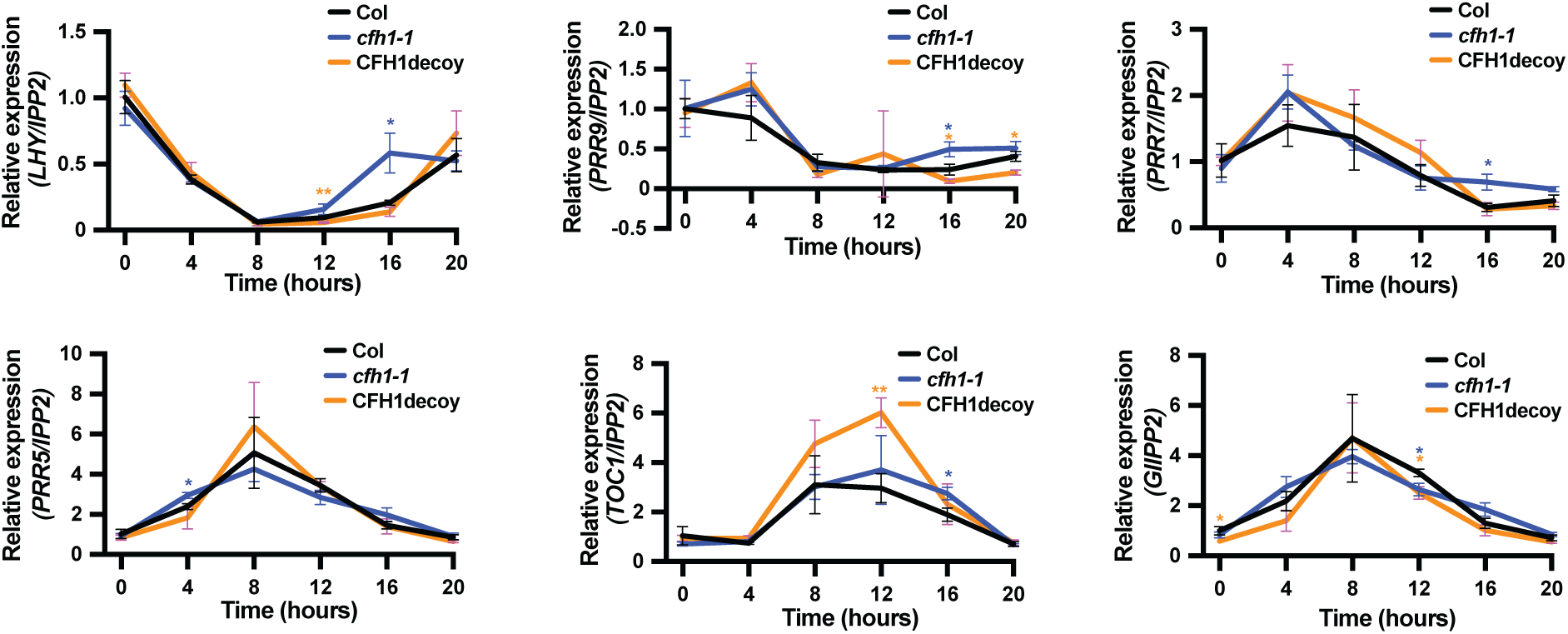
Expression of clock genes in *cfh1-1* mutant and *CFH1* decoy. qRT-PCR of *LHY*, *PRR9, PRR7, PRR5, TOC1,* and *GI* in wild-type (Col), *cfh1-1* mutant, and *CFH1* decoy plants grown in constant white light. *IPP2* was used as an internal control. Error bars indicate SD (n = 3 replicates). *, p ≤ 0.05; **, p ≤ 0.01 (Welch’s t-test). Color of the star indicates the statistical analysis between the colored genotype (blue, *cfh1-1*; orange, *CFH1* decoy) and wild-type (Col).

**Fig. S5.**
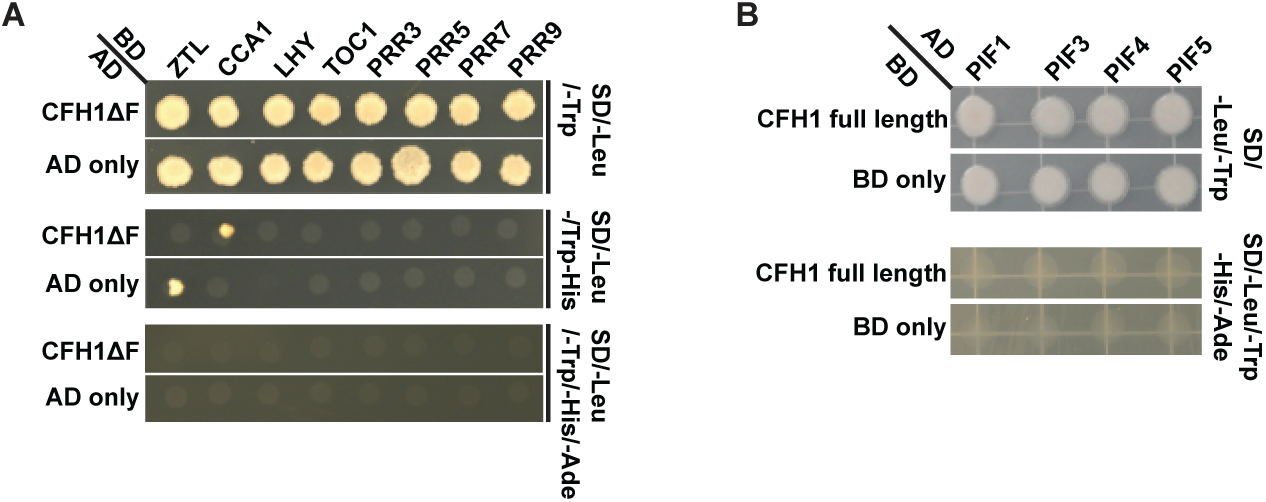
Yeast two-hybrid assay. (**A**). Yeast transformed with GAL4 DNA-binding domain (BD) fused to ZTL, CCA1, LHY, TOC1, PRR3, PRR5, PRR7, PRR9 and GAL4 activation domain (AD) fused to CFH1ΔF (BD-CFH1ΔF) were grown in SD-Leu-Trp medium for autotrophic selection and on SD-Leu-Trp-His or SD-Leu-Trp-His-Ade medium to test for interactions. (**B**). Yeast transformed with GAL4 DNA-binding domain (BD) fused to CFH1 full length and GAL4 activation domain (AD) fused to PIF1, PIF3, PIF4, and PIF5 were grown in SD-Leu-Trp medium for autotrophic selection and on SD-Leu-Trp-His-Ade medium to test for interactions.

**Fig. S6.**
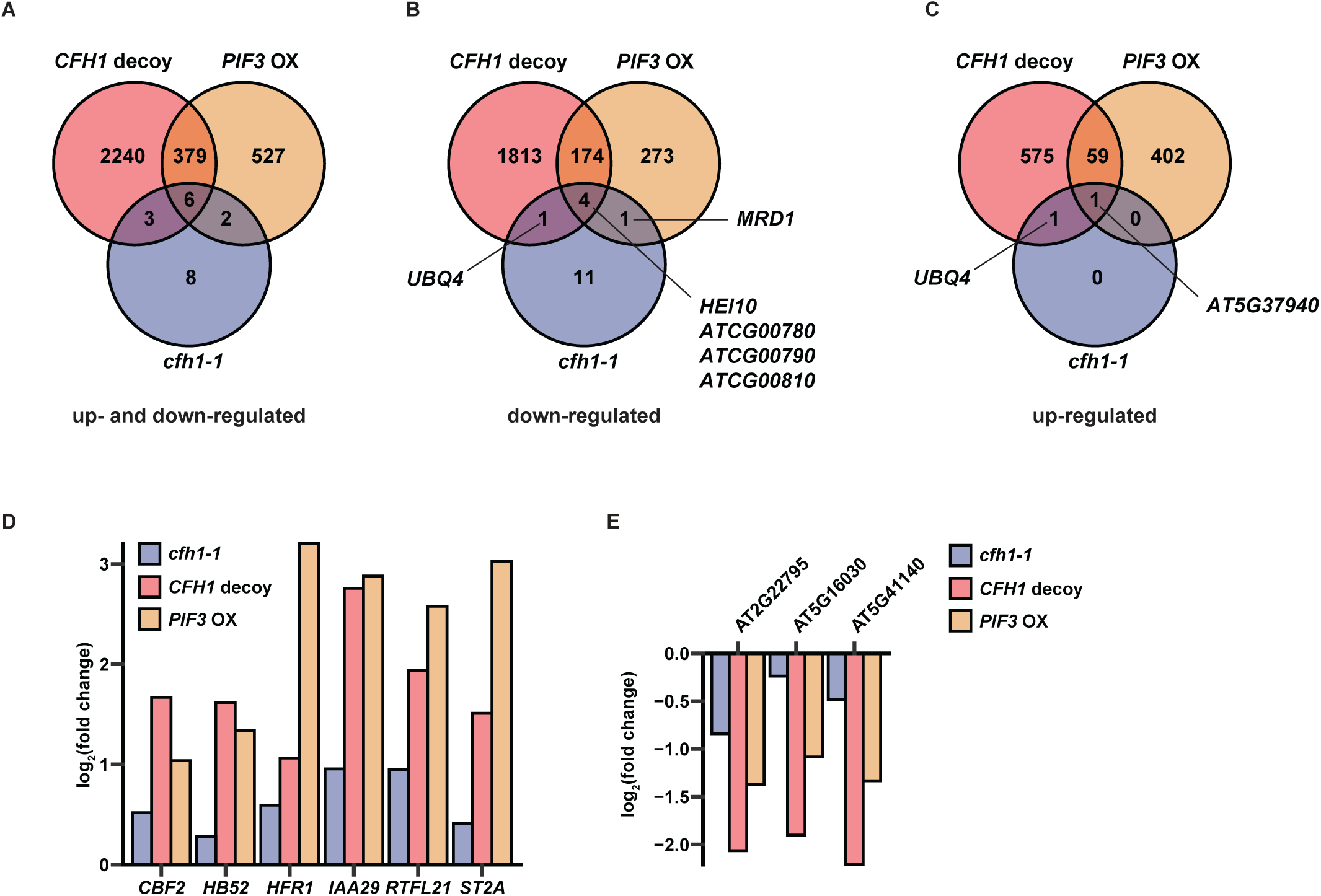
Differentially expressed genes in *CFH1* decoy, PIF3 OX, and *cfh1-1* mutant as compared to wild-type (Col). (**A-C**). Venn diagrams depicting the number of (**A**) all differentially expressed genes, (**B**) down-regulated genes, (**C**) up-regulated genes, in *cfh1-1* mutant, *CFH1* decoy, and *PIF3* OX transgenic lines as compared to the wild type (Col). (**D-E**). Expression fold change of (**D**) six co-induced PIF3-bound genes and (**E**) three co-repressed PIF3-bound genes (*37*) in *cfh1-1* mutant, *CFH1* decoy, *PIF3* OX transgenic lines as compared to the wild type (Col).

**Fig. S7.**
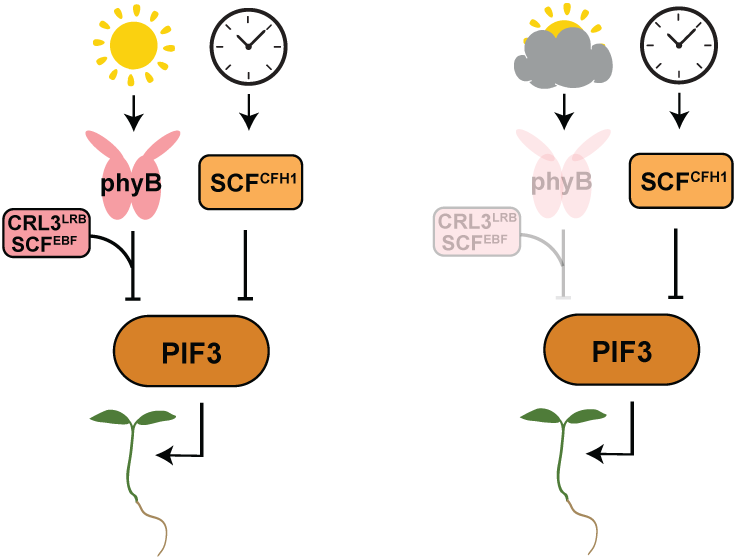
Proposed model of CFH1 function. The circadian clock phases CFH1 expression to the early morning, which in turn mediates the stability of PIF3 protein to control hypocotyl length. This regulatory mechanism functions alongside another pathway involving PhyB-LRB/EBF mediated stabilization of PIF3 protein. In conditions where red light signaling is compromised, the circadian clock ensures proper hypocotyl elongation through CFH1 mediated degradation of PIF3.

## References

1. P. Mas, W. Y. Kim, D. E. Somers, S. A. Kay, Targeted degradation of TOC1 by ZTL modulates circadian function in Arabidopsis thaliana. Nature 426, 567–570 (2003).

2. B. Grima et al., The F-box protein slimb controls the levels of clock proteins period and timeless. Nature 420, 178–182 (2002).

3. H. W. Ko, J. Jiang, I. Edery, Role for Slimb in the degradation of Drosophila Period protein phosphorylated by Doubletime. Nature 420, 673–678 (2002).

4. J. Lee et al., The E3 ubiquitin ligase adaptor Tango10 links the core circadian clock to neuropeptide and behavioral rhythms. Proc Natl Acad Sci U S A 118, (2021).

5. Y. H. Song, R. W. Smith, B. J. To, A. J. Millar, T. Imaizumi, FKF1 conveys timing information for CONSTANS stabilization in photoperiodic flowering. Science 336, 1045–1049 (2012).

6. T. C. Mockler et al., The DIURNAL project: DIURNAL and circadian expression profiling, model-based pattern matching, and promoter analysis. Cold Spring Harb Symp Quant Biol 72, 353–363 (2007).

7. W. Liu et al., A metabolic daylength measurement system mediates winter photoperiodism in plants. Dev Cell 56, 2501–2515 e2505 (2021).

8. D. C. Nelson, J. Lasswell, L. E. Rogg, M. A. Cohen, B. Bartel, FKF1, a clock-controlled gene that regulates the transition to flowering in Arabidopsis. Cell 101, 331–340 (2000).

9. J. M. Gendron, C. C. Leung, W. Liu, Energy as a seasonal signal for growth and reproduction. Curr Opin Plant Biol 63, 102092 (2021).

10. J. M. Gendron, D. Staiger, New Horizons in Plant Photoperiodism. Annu Rev Plant Biol 74, 481–509 (2023).

11. C. M. Lee et al., Decoys Untangle Complicated Redundancy and Reveal Targets of Circadian Clock F-Box Proteins. Plant physiology 177, 1170–1186 (2018).

12. A. Feke et al., Decoys provide a scalable platform for the identification of plant E3 ubiquitin ligases that regulate circadian function. Elife 8, (2019).

13. C. M. Lee et al., GIGANTEA recruits the UBP12 and UBP13 deubiquitylases to regulate accumulation of the ZTL photoreceptor complex. Nat Commun 10, 3750 (2019).

14. A. M. Feke, J. Hong, W. Liu, J. M. Gendron, A Decoy Library Uncovers U-Box E3 Ubiquitin Ligases That Regulate Flowering Time in Arabidopsis. Genetics 215, 699–712 (2020).

15. C. M. Lee, C. Adamchek, A. Feke, D. A. Nusinow, J. M. Gendron, Mapping Protein-Protein Interactions Using Affinity Purification and Mass Spectrometry. Methods Mol Biol 1610, 231–249 (2017).

16. K. Ao et al., SCF(SNIPER7) controls protein turnover of unfoldase CDC48A to promote plant immunity. New Phytol 229, 2795–2811 (2021).

17. M. Hersch et al., Light intensity modulates the regulatory network of the shade avoidance response in Arabidopsis. Proc Natl Acad Sci U S A 111, 6515–6520 (2014).

18. T. Zielinski, A. M. Moore, E. Troup, K. J. Halliday, A. J. Millar, Strengths and limitations of period estimation methods for circadian data. PLoS One 9, e96462 (2014).

19. N. Dalchau et al., The circadian oscillator gene GIGANTEA mediates a long-term response of the Arabidopsis thaliana circadian clock to sucrose. Proc Natl Acad Sci U S A 108, 5104–5109 (2011).

20. J. L. Pruneda-Paz, A functional genomics approach reveals CHE as a component of the Arabidopsis circadian clock (vol 325, pg 1481, 2009). Science 326, 366–366 (2009).

21. J. M. Gendron et al., Arabidopsis circadian clock protein, TOC1, is a DNA-binding transcription factor. Proc Natl Acad Sci U S A 109, 3167–3172 (2012).

22. M. Chen et al., Arabidopsis HEMERA/pTAC12 initiates photomorphogenesis by phytochromes. Cell 141, 1230–1240 (2010).

23. C. Y. Yoo et al., Direct photoresponsive inhibition of a p53-like transcription activation domain in PIF3 by Arabidopsis phytochrome B. Nat Commun 12, 5614 (2021).

24. P. Leivar et al., The Arabidopsis phytochrome-interacting factor PIF7, together with PIF3 and PIF4, regulates responses to prolonged red light by modulating phyB levels. Plant Cell 20, 337–352 (2008).

25. J. Dong et al., Light-Dependent Degradation of PIF3 by SCF(EBF1/2) Promotes a Photomorphogenic Response in Arabidopsis. Curr Biol 27, 2420–2430 e2426 (2017).

26. W. Ni et al., A mutually assured destruction mechanism attenuates light signaling in Arabidopsis. Science 344, 1160–1164 (2014).

27. J. S. O’Neill et al., Circadian rhythms persist without transcription in a eukaryote. Nature 469, 554–558 (2011).

28. D. Ezer et al., The evening complex coordinates environmental and endogenous signals in Arabidopsis. Nat Plants 3, 17087 (2017).

29. Y. Zhang et al., Central clock components modulate plant shade avoidance by directly repressing transcriptional activation activity of PIF proteins. Proc Natl Acad Sci U S A, 3261–3269 (2020).

30. J. Y. Zhu, E. Oh, T. Wang, Z. Y. Wang, TOC1-PIF4 interaction mediates the circadian gating of thermoresponsive growth in Arabidopsis. Nat Commun 7, 13692 (2016).

31. J. Soy et al., Phytochrome-imposed oscillations in PIF3 protein abundance regulate hypocotyl growth under diurnal light/dark conditions in Arabidopsis. Plant J 71, 390–401 (2012).

32. C. Y. Yoo et al., Phytochrome activates the plastid-encoded RNA polymerase for chloroplast biogenesis via nucleus-to-plastid signaling. Nat Commun 10, 2629 (2019).

33. X. Liu et al., PHYTOCHROME INTERACTING FACTOR3 associates with the histone deacetylase HDA15 in repression of chlorophyll biosynthesis and photosynthesis in etiolated Arabidopsis seedlings. Plant Cell 25, 1258–1273 (2013).

34. A. Oda, S. Fujiwara, H. Kamada, G. Coupland, T. Mizoguchi, Antisense suppression of the Arabidopsis PIF3 gene does not affect circadian rhythms but causes early flowering and increases FT expression. FEBS Lett 557, 259–264 (2004).

35. B. C. Jiang et al., PIF3 is a negative regulator of the CBF pathway and freezing tolerance in Arabidopsis. P Natl Acad Sci USA 114, E6695–E6702 (2017).

36. A. Feke, M. Vanderwall, W. Liu, J. M. Gendron, Functional domain studies uncover novel roles for the ZTL Kelch repeat domain in clock function. PLoS One 16, e0235938 (2021).

37. A. Pfeiffer, H. Shi, J. M. Tepperman, Y. Zhang, P. H. Quail, Combinatorial Complexity in a Transcriptionally Centered Signaling Hub in Arabidopsis. Molecular Plant 7, 1598–1618 (2014).

38. D. A. Nusinow et al., The ELF4-ELF3-LUX complex links the circadian clock to diurnal control of hypocotyl growth. Nature 475, 398–402 (2011).

39. J. Dong et al., Arabidopsis DE-ETIOLATED1 represses photomorphogenesis by positively regulating phytochrome-interacting factors in the dark. Plant Cell 26, 3630–3645 (2014).

40. H. Huang et al., Identification of Evening Complex Associated Proteins in Arabidopsis by Affinity Purification and Mass Spectrometry. Mol Cell Proteomics 15, 201–217 (2016).

41. M. D. Curtis, U. Grossniklaus, A gateway cloning vector set for high-throughput functional analysis of genes in planta. Plant Physiol 133, 462–469 (2003).

42. T. Nakagawa et al., Development of series of gateway binary vectors, pGWBs, for realizing efficient construction of fusion genes for plant transformation. J Biosci Bioeng 104, 34–41 (2007).

43. C. LeBlanc et al., Increased efficiency of targeted mutagenesis by CRISPR/Cas9 in plants using heat stress. Plant J 93, 377–386 (2018).

44. G. Yu et al., A bacterial effector protein prevents MAPK-mediated phosphorylation of SGT1 to suppress plant immunity. PLoS Pathog 16, e1008933 (2020).

45. Y. Qiu et al., HEMERA Couples the Proteolysis and Transcriptional Activity of PHYTOCHROME INTERACTING FACTORs in Arabidopsis Photomorphogenesis. Plant Cell 27, 1409–1427 (2015).

46. X. Wang et al., CSN1 N-terminal-dependent activity is required for Arabidopsis development but not for Rub1/Nedd8 deconjugation of cullins: a structure-function study of CSN1 subunit of COP9 signalosome. Mol Biol Cell 13, 646–655 (2002).

47. C. C. Leung, D. A. Tarte, L. S. Oliver, Q. Wang, J. M. Gendron, Systematic characterization of photoperiodic gene expression patterns reveals diverse seasonal transcriptional systems in Arabidopsis. PLoS Biol 21, e3002283 (2023).

48. A. M. Bolger, M. Lohse, B. Usadel, Trimmomatic: a flexible trimmer for Illumina sequence data. Bioinformatics 30, 2114–2120 (2014).

49. R. Patro, G. Duggal, M. I. Love, R. A. Irizarry, C. Kingsford, Salmon provides fast and bias-aware quantification of transcript expression. Nat Methods 14, 417–419 (2017).

50. C. Soneson, M. I. Love, M. D. Robinson, Differential analyses for RNA-seq: transcript-level estimates improve gene-level inferences. F1000Res 4, 1521 (2015).

51. Y. Zhang et al., A quartet of PIF bHLH factors provides a transcriptionally centered signaling hub that regulates seedling morphogenesis through differential expression-patterning of shared target genes in Arabidopsis. PLoS Genet 9, e1003244 (2013).

52. Z. Gu, R. Eils, M. Schlesner, Complex heatmaps reveal patterns and correlations in multidimensional genomic data. Bioinformatics 32, 2847–2849 (2016).

